# Platelet-neutrophil aggregates promote skin pathology in psoriasis

**DOI:** 10.1101/526236

**Authors:** Franziska Herster, Zsofia Bittner, Marius Cosmin Cordrea, Nate Archer, Martin Heister, Markus W. Löffler, Simon Heumos, Joanna Wegner, Ramona Businger, Michael Schindler, David Stegner, Knut Schäkel, Stephan Grabbe, Kamran Ghoreschi, Lloyd Miller, Alexander N.R. Weber

**Author notes:** Contact information (Corresponding Author and Lead Contact) Alexander N. R. Weber, Interfaculty Institute for Cell Biology, Department of Immunology, University of Tübingen, Auf der Morgenstelle 15, 72076 Tübingen, Germany.

## Abstract

Psoriasis is a frequent systemic inflammatory autoimmune disease characterized primarily by skin lesions with massive infiltration of leukocytes but frequently also presents with cardiovascular comorbidities. Especially polymorphonuclear neutrophils (PMNs) abundantly infiltrate psoriatic skin but the cues that prompt PMNs to home to the skin are not well defined. To identify PMN surface receptors that may explain PMN skin homing in psoriasis patients, we screened 332 surface antigens on primary human blood PMNs from healthy donors and psoriasis patients. We identified platelet surface antigens as a defining feature of psoriasis PMNs, due to a significantly increased aggregation of neutrophils and platelets in the blood of psoriasis patients. Similarly, in the imiquimod-induced experimental in vivo model of psoriasis, disease induction promoted PMN-platelet aggregate formation. In psoriasis patients, disease directly correlated with blood platelet counts and platelets were detected in direct contact with PMNs in psoriatic but not healthy skin. Importantly, depletion of circulating platelets in vivo ameliorated disease severity significantly, indicating that the intimate relationship of PMNs and platelets may be relevant for psoriasis pathology and disease severity, and potentially for psoriasis-associated cardiovascular comorbidities.

**Key points:**

- Human neutrophils in psoriasis patient blood show a distinct ‘platelet signature’ of surface antigens
- Platelets congregate with neutrophils in psoriatic skin lesions
- Circulating platelets contribute to psoriasis skin pathology

## Introduction

Psoriasis is a frequent, chronic, immune-mediated inflammatory skin disease of unknown aetiology (Eberle et al., 2016). Its most common form, plaque psoriasis, shows epidermal hyperplasia, increased endothelial proliferation and a prominent infiltrate of polymorphonuclear neutrophils (PMNs) (Eberle et al., 2016;Schon et al., 2017). The accumulation of PMNs in psoriatic plaques and micro-abscesses is accompanied by an increase of PMNs in the circulation of psoriasis patients but their precise role in the disease remains enigmatic (Sen et al., 2013),(Schon et al., 2017). Furthermore, it remains unclear which factors prompt PMNs, plasmacytoid dendritic cells and T cells to accumulate in psoriatic skin. The latter cells drive a chronic phase of disease, dominated by IL-17 cytokines (Eberle et al., 2016;Schon et al., 2017;Schon and Erpenbeck, 2018). Apart from its strong manifestation in the skin, psoriasis is now considered a systemic disease, and besides frequent joint involvement (psoriasis arthritis), alterations in circulating immune cell subsets have been reported: for example, the frequency CD16^+^ monocytes is altered in psoriasis patients and these cells were observed to aggregate more with other monocytes or lymphocytes in patient blood (Golden et al., 2015). Changes within the T cell compartment have also been reported (Langewouters et al., 2008). Additionally, a strong link between psoriasis and cardiovascular comorbidities has been noted (Vena et al., 2010;Boehncke, 2018;Caiazzo et al., 2018), for which IL-17A may be an important link (Schuler et al., 2018). Regarding the cardiovascular events, psoriasis severity correlates with the incidence cerebrovascular, peripheral vascular and heart structural disorders (reviewed in (Vena et al., 2010)). Here a potential etiology involving platelets has been proposed (Tamagawa-Mineoka, 2015) but not been proven experimentally, especially in terms of a direct link between PMNs and platelets in humans.

Recent research has uncovered an intimate relationship between different leukocyte populations, including PMNs, and platelets. Interestingly, the existence of direct leukocyte-platelet aggregates in vitro and in human blood has been known for some time (Jungi et al., 1986;de Bruijne-Admiraal et al., 1992). Ludwig et al. reported that PSGL-1-P-selectin-mediated interactions between platelets and leukocytes promoted rolling in murine skin micro vessels and the same receptors were responsible for activated platelets to interact with murine PBMCs when co-cultured in vitro (Ludwig et al., 2004). When these co-cultured PBMCs were infused in mice, rolling in the murine skin microvasculature was observed. The authors speculated whether platelets might also prompt leukocyte invasion into the inflamed skin and noted that P-selectin expression correlated with psoriasis score. However, their studies were largely based on transfusion experiments and did not focus on PMNs. More recently, the capacity of platelets to direct PMN extravasation experimental was extensively studied in in vivo models (Sreeramkumar et al., 2014), and (Zuchtriegel et al., 2016) proposed that initially CD40-CD40L interactions mediate leukocyte capture at the vessel wall for PSGL-1-P-selectin interactions to guide subsequent extravasation. Whilst this is required for peripheral defense, in rheumatoid arthritis it was also observed that platelets attract neutrophils into the synovium where they become trapped and contribute to disease severity (Habets et al., 2013). Whether PMN-platelet aggregates occur in human psoriasis patients and whether they contribute to psoriasis skin inflammation was not experimentally studied.

In order to identify surface antigens on human peripheral blood immune cells that might be involved in their skin-homing in psoriasis, we screened 322 surface antigens in PMNs, monocytes and T- and B-lymphocytes from psoriasis patients and healthy controls. This unbiased approach identified surface antigen signatures specific for different blood immune cell populations in psoriasis. For PMNs platelet markers were significantly increased and this surface antigen signature was attributable to direct PMN-platelet aggregates. Such aggregates were also observed upon psoriasis induction in mice and in the skin of psoriasis patients. Interestingly, depletion of platelets in the imiquimod experimental in vivo setting of psoriasis drastically decreased ear and epidermal thickness of the skin. Collectively, our results establish a functional link between circulating PMN-platelet aggregates and disease activity in the psoriatic skin.

## Materials and Methods

### Reagents

All chemicals were from Sigma or Invitrogen, respectively, unless otherwise stated in Supplementary materials. Antibodies and recombinant cytokines are listed in Table S1.

### Mice

Sex-and age-matched C57BI/6J SPF mice (Jackson Laboratories) bred and maintained according to local animal welfare guidelines and applicable regulations (see Supplemental information) were used at the age of 6-8 weeks.

### Study participants and sample acquisition

All patients and healthy blood donors included in this study provided their written informed consent before study participation. Approval for use of their biomaterials was obtained by the local ethics committees at the University Hospitals of Tübingen and Heidelberg, in accordance with the principles laid down in the Declaration of Helsinki as well as applicable laws and regulations. All blood or skin samples obtained from psoriasis patients (median age 41.8 years, psoriasis area severity index (PASI) ≥10 (except for 2 donors in Figs. 3B,C and S3B,C where PASI was >4.5) no systemic treatments at the time of blood/skin sampling) were obtained at the University Hospitals Tübingen or Heidelberg, Departments of Dermatology, and were processed simultaneously with samples from at least one healthy donor matched for age and sex (recruited at the University of Tübingen, Department of Immunology). Skin sections were obtained from 11 patients with plaque psoriasis and 1 patient with psoriasis guttata. Platelet counts were determined in the course of established clinical routines at the time of study blood sampling.

### Cell surface marker expression screening in whole blood

A cell surface antigen screening was performed using the LegendScreen from Biolegend (Fig. 1A). Whole blood (EDTA) was drawn from 5 psoriasis patients and 5 sex- and age-matched controls. Erythrocyte lysis was performed for 5 min at 4°C on a roller shaker. After a short spin, FC block was performed and the cells were stained with anti-CD3, -CD15 and -CD19, excluding dead cells using Zombie Yellow. Subsequently, the stained cells were aliquoted into 96 well, each containing a PE-labeled antibody directed against one of 332 surface antigens, and 13 isotype controls in PE. The following washing and further steps were performed using the manufacturer’s instructions, except that one kit was divided for the measurement of four donors. FACS measurements were performed using a MACSQuant analyzer (Miltenyi) and subsequently FlowJo V10 was used to analyze the data. The gating strategy is depicted in Fig. S1 and T cells, PMNs and B cells gated according to the Abs in the master mix. Monocytes were gated by granularity and size but not additionally verified with CD14 staining. However, in the well containing anti-CD14-PE Abs, all gated events were CD14-positive.

**Figure 1:**
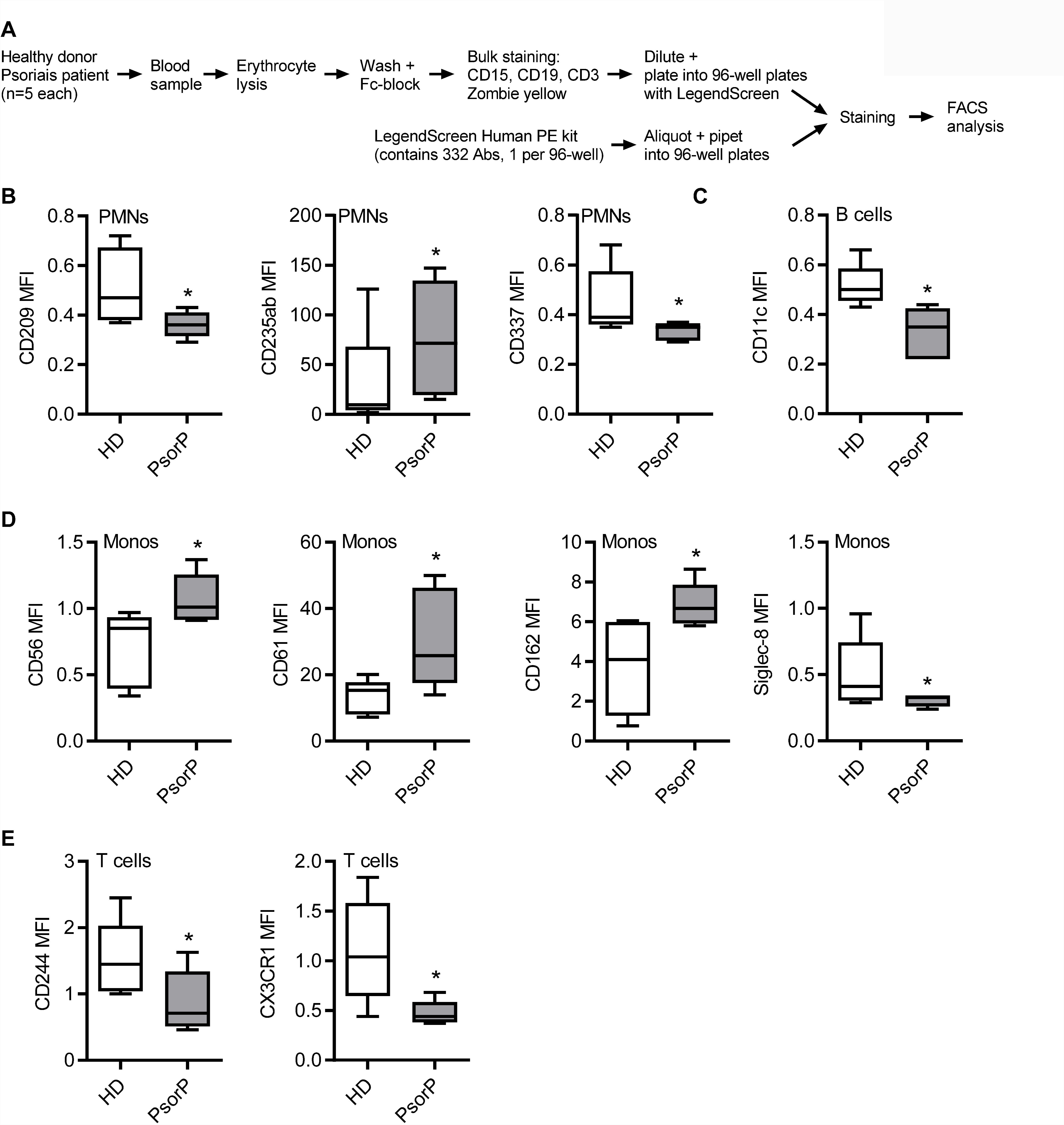
Immune cells in psoriasis show distinct surface antigens. (A) Schematic overview of surface antigen screen procedure. (B-E) Selected surface antigens with significant differences in MFI between healthy donors (HDs) and psoriasis patients (PsorP), n=5 each. B-E represent combined data (mean+SD) from ‘n’ biological replicates. * p<0.1 nominal by two-way ANOVA followed by Tukey’s multiple comparisons correction.

### Differential expression analysis of surface marker screening data

Conceptually, the goal was to identify surface antigens which are significantly different between patients and healthy donors within the targeted cell types. This relationship was formulated as MFI ∼ health_status + cell_type, where the mean fluorescence intensity (MFI) depends on the two main factors: health_status with levels {patient, healthy donor} and cell_type with levels (Zuchtriegel et al., 2016 T cells), gated as indicated in Fig. S1A. All cell types were measured simultaneously in one FACS screen per subject (patient or healthy donor) and the resulting intrinsic within-subject variance (across cell types) was accounted for by extending MFI ∼ health_status + cell_type + subject/cell_type. In this notation, cell_type is “nested” within “subject”, accommodating the above. The implementation of the analysis was done in R [version 3.4.4, with linear mixed models using the R package nlme (version 3.1-131.1, details and code upon request, Refs. (Bates et al., 2000;Bates et al., 2015)). The fitted models were subject to post-hoc analysis with Tukey’s “Honest Significant Difference” test to compute adjusted pair-wise differences among the cell types (Bretz et al.). The Ismeans (version 2.27-61) (Lenth, 2016) R package implementation was used in order to compute the adjustments. With Ismeans’ pairs all pair-wise contrasts of patient versus healthy donor by the given cell_type were calculated. From all contrasts of the different cell_type levels the p-values and the fold changes of all surface antigens were extracted. This analysis provides multiple-comparison adjusted p-values and fold changes between patients versus healthy donors for each cell type. For the subsequent platelet analysis an unpaired t-test method was used as only one cell type had to be considered. Receptors with MFI values missing for more than 1 donor per group were filtered out. For the generation of the PCA plots, the R package ggplot2 (version 2.2.1) was used.

### Flow cytometry

200 μl of the cell suspension was transferred into a 96 well round bottom plate and spun down for 5 min at 322 × g, 4 °C. FcR block was performed using pooled human serum diluted 1:10 in FACS buffer (PBS, 1 mM EDTA, 2% FBS heat inactivated) for 15 min at 4 °C. After washing, the samples were stained for 20-30 min at 4°C in the dark. Thereafter, fixation buffer (4% PFA in PBS) was added to the cell pellets and incubated for 10 min at RT in the dark. After an additional washing step, the cell pellets were resuspended in 100 μl FACS buffer. Measurements were performed on a MACSQuant analyzer (Miltenyi). Analysis was performed using FlowJo V10 analysis software.

### Fluorescence Microscopy of fixed whole blood cells

The cells were seeded in a 96 well plate 200 μl cell suspension per well. FcR block, staining, fixation and permeabilization were performed as for flow cytometry. After 0.05% Saponin permeabilization, nuclear DNA was stained using Hoechst 33342 (1 μg/ml, Thermo Fisher). The cell pellets were resuspended in 50-100 μl FACS buffer. 40 μl of the cell suspension was pipetted on a Poly-L-Lysine coated coverslip (734-1005, Corning) and the cells were left to attach for one hour in the dark. ProLong Diamond Antifade (P36965, Thermo Fisher) was used to mount the coverslips on uncoated microscopy slides. The slides were left to dry overnight at RT in the dark and were then stored at 4 °C before microscopy. The measurements were conducted with a Nikon Ti2 eclipse (100x magnification) and the analysis was performed using ImageJ/Fiji analysis software (Schindelin et al., 2012).

### Fluorescence Microscopy of tissue samples (human and mouse)

Skin samples from psoriasis patients with a PASI score ≥10 and without systemic treatment at the time of sampling, and healthy skin samples and skin samples from IMQ treated mice, were paraffin-embedded according to standard procedures were deparaffinized and rehydrated using Roti Histol (Roth, 6640.1) and decreasing concentrations of ethanol (100%, 95%, 80% and 70%). After rinsing in ddH_2_O, antigen retrieval was performed by boiling for 10-20 min in citrate buffer (0.1 M, pH=6). The skin tissue was then washed 3 times for 5 min with PBS. Blocking was performed using pooled human serum (1:10 in PBS) for 30 min at RT. The primary antibody was added either overnight at 4°C or for 1 hour at RT. After 3 washes, the secondary antibody was incubated for 30 min at RT in the dark. Thereafter, the samples were washed again and Hoechst 33342 (1 μg/ml, ThermoFisher) was added for 5 min. Then 3 last washes were performed before using ProLong Diamond Antifade (P36965, Thermo Fisher) for mounting. The samples were left to dry overnight at RT in the dark before being used for microscopy or stored at 4°C. The specimens were analyzed on a Nikon Ti2 eclipse microscope (40x-60x magnification) and the analysis was performed using ImageJ/Fiji analysis software.

### Platelet analysis in the imiquimod model of psoriatic skin inflammation

Prior and after daily topical application of 5% imiquimod cream to the ears of anesthetized (2% isoflurane) mice ear thickness was measured with a manual caliper (Peacock) on d0 to d5. For FACS analysis, retro-orbital blood samples were collected for FACS analysis on d0 and d5, diluted in TBS containing 5 U/ml Heparin and subsequently PBS and an aliquot stained using anti-Ly6C, anti-CD45, anti-CD41, anti-Ly6G, anti-CD11b, propidium iodide and TruStain fcX (see Table S1). Full thickness ear skin was excised on d5, fixed in 10% formalin and paraffin-embedded. Skin cross-sections (4 μm) were stained by H&E according to standard procedures. At least 10 epidermal thickness measurements per mouse were averaged. For details see Supplemental information.

### Platelet depletion protocol

For platelet depletion in mice, 4 μg/g of anti-CD42b (clone R300, Emfret Analytics) or rat IgG isotype control (clone R301, Emfret Analytics) in sterile PBS was administered i.v. one day before, and 2 μg/g administered i.p. 3 days after the first imiquimod treatment.

### Statistics

Statistics regarding differential surface antigen expression analysis are described above. All other experimental data were analyzed using Excel 2010 (Microsoft) and/or GraphPad Prism 6, 7 or 8, microscopy data with ImageJ/Fiji, flow cytometry data with FlowJo V10. In case of extreme values, outliers were statistically identified using the ROUT method at high stringency (0.5%) and normality tested using the Shapiro-Wilk test for the subsequent choice of a parametric or non-parametric test, p-values (α =0.05) were then calculated and multiple testing was corrected for in Prism, as indicated in the figure legends. Values < 0.05 were considered statistically as significant and denoted by * throughout.

## Results

### Circulating neutrophils in psoriasis show a distinct platelet signature of surface antigens

To phenotype immune cells in psoriasis patients with a view to identifying surface antigens disparately regulated in psoriasis PMNs, we combined the so-called LegendScreen, a screening format consisting of antibodies directed against 332 different surface antigens and 10 corresponding isotype controls, with a whole blood staining assay using anti-CD3, CD15, CD19 Abs and a live/dead marker (Fig. 1A), based on previously published work (Graessel et al., 2015). Five psoriasis patients with PASI ≥10 and without systemic therapy (at the time of blood drawing) and 5 healthy donors were analyzed, the major cell populations gated as described in Fig. S1A) and mean fluorescence intensities (MFIs) acquired for each of the 322 surface antigens (Table SI). Differential expression analysis was performed by two-way ANOVA followed by Tukey’s multiple comparisons correction (see Methods and Supplemental R code). Conceptually, the goal of the differential expression analysis was to identify surface antigens which are significantly different between patients and healthy donors within the targeted cell types. Figs. 1B-E and S2A-C show selected surface antigens with significantly different MFIs between patient and control PMNs, FSC-SSC-gated monocytes (see methods), CD3 T cells and CD19 B cells. Evidently, >30 surface antigens were significantly altered across the four major immune cell populations, with monocytes being most strongly affected (Table S2).

Although individual surface markers were of interest, we next considered whether a certain combination (‘signature’) of molecules was suitable to discriminate patients and healthy donors. In order to check whether the most differently expressed surface antigens (based on a nominal p-value <0.1) were able to separate the two groups a principal component analysis (PCA) was performed. (Fig. 2A-D). This showed that especially for PMNs (Fig. 2A), psoriasis patient samples clustered together tightly. The antigens contributing most to the separation of both groups were CD41, CD61 and CD235ab which are linked more closely to patient data points (Fig. 2A). For monocytes, psoriasis patient samples clustered less tightly and a greater number of markers contributed to group separation (Fig. 2B). B cells also showed only a moderate clustering and all markers with discriminatory power were lower in patients than healthy donors (Fig. 2C, *cf.* also Fig. S1D). For T cells, patient and healthy donor samples were only loosely clustered (Fig. 2D). Collectively, our analysis in untreated whole blood samples shows significant differences in surface antigen expression across circulating immune cell populations and that selected groups of surface antigens are able to discriminate patients and healthy donors on the basis of PMN surface antigens.

**Figure 2:**
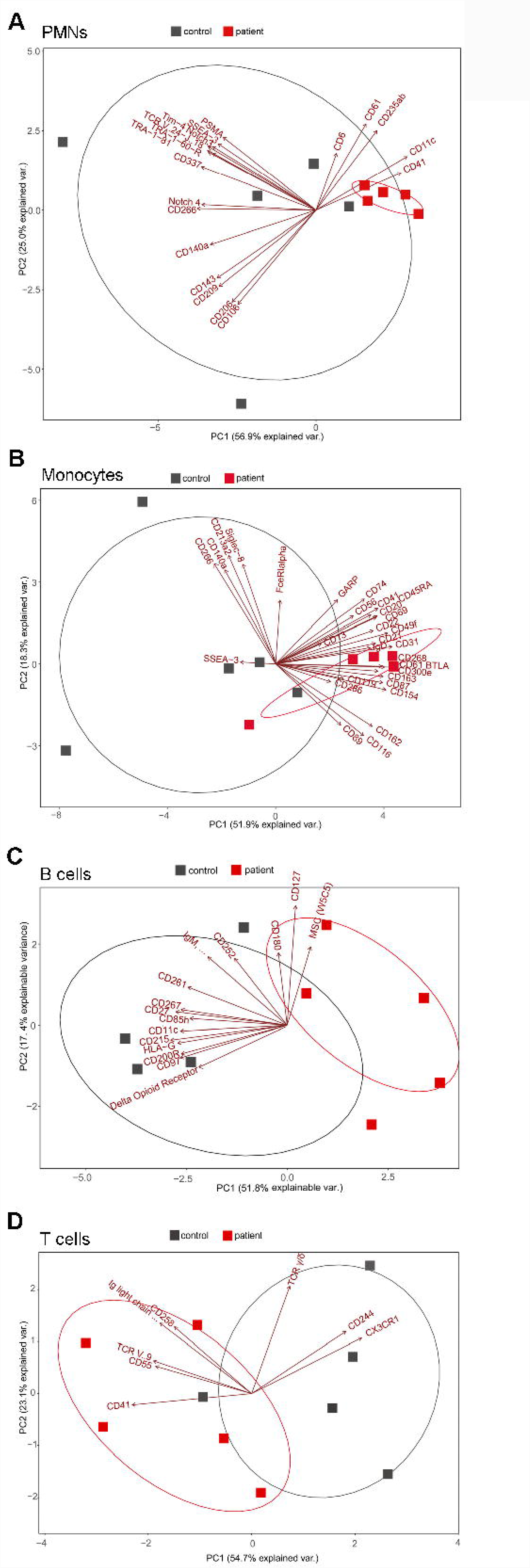
Psoriasis PMNs show a distinct ‘platelet signature’. (A-D) Principal component analyses of PMN (A), B cell (B), monocyte (C) or T cell (D) surface antigens for healthy donors (red) and psoriasis patients (blue) with top significant antigens (based on nominal p-value <0.1, n=5 each) contributing to a separation of patients and healthy donors. A-D represent combined data (mean+SD) from ‘n’ biological replicates (each dot represents one donor).

**Figure 3:**
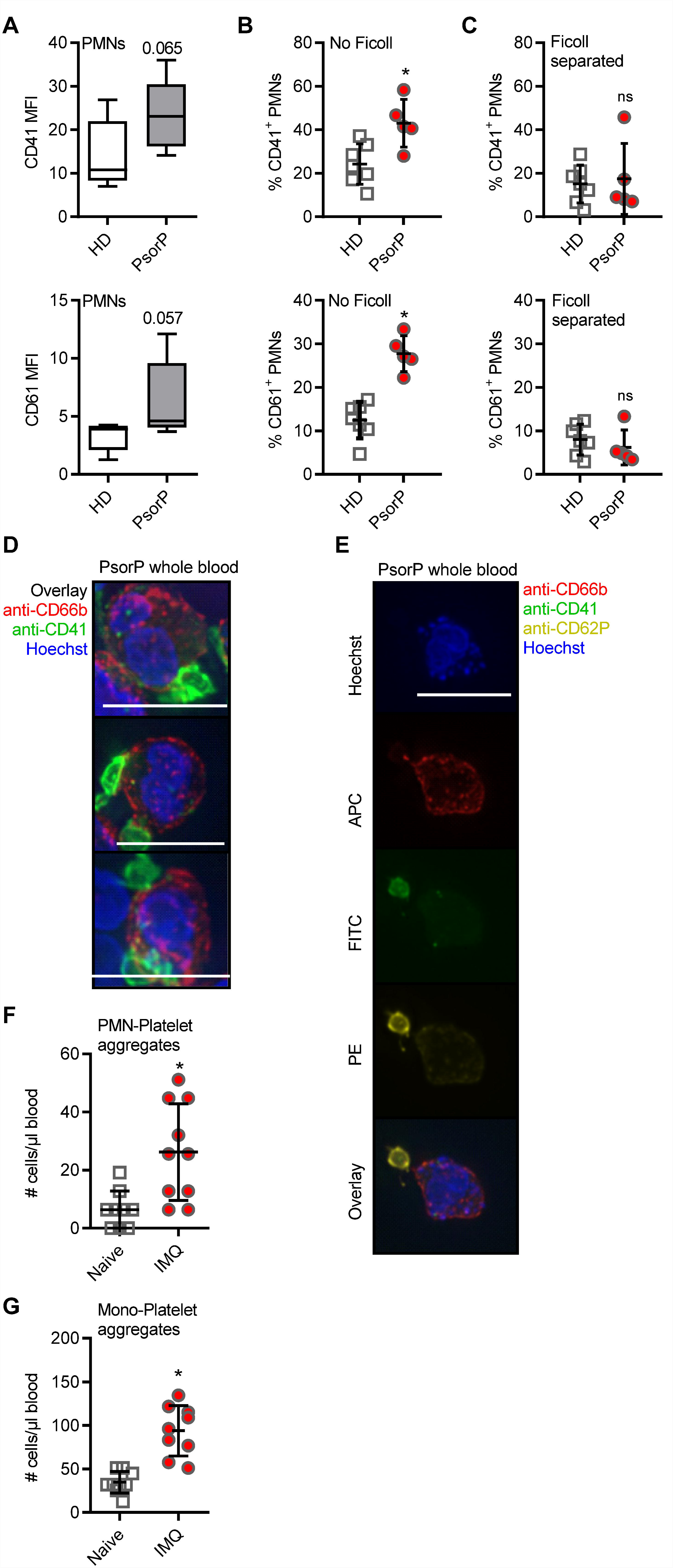
Psoriasis PMNs directly interact with platelets more prominent in psoriasis blood samples and upon in vivo psoriasis induction. (A) Expression analysis screening results for CD41 (upper) and CD61 (lower panel) MFIs on gated PMNs between healthy donors (HDs) and psoriasis patients (PsorP), n=5 each. (B+C) Flow cytometric analysis of CD41- or CD61-positive PMNs (defined as CD15^+^CD66b^+^and CD41^+^ or CD61^+^) analyzed in HD or PsorP (HD n=7, PsorP n=5) in whole blood samples (B) or Ficoll density gradient centrifugation. (D) Fluorescence microscopy of PMNs in a PsorP whole blood stained as indicated (scale bar = 10 μm). (E) as in D. (F and G) Mean number of PMN-platelet (F) or PMN-monocyte (G) aggregates comparing naive (day 0) and IMQ-treated (day 5) isotype mice (n=10 each). A-C and F-G represent combined data (mean+SD) from ‘n’ biological replicates (in B, C, F and G each dot represents one donor or mouse). In D, E one representative of ‘n’ biological replicates (donors) is shown (mean+SD). * p<0.05 according to two-way ANOVA followed by Tukey’s multiple comparisons correction (A), Mann-Whitney test (B, C), unpaired Student’s t-test (F, G).

### Circulating neutrophils in psoriasis directly interact with platelets

Since PMN infiltration is a hallmark of psoriasis we focused on PMN surface antigens and noted that several of the aforementioned psoriasis-associated surface antigens, e.g. CD41 and CD61, are typically associated with platelets. These data indicated a ‘platelet-like’ signature on psoriasis PMNs, and, upon re-inspection of the screen data, CD41 and CD61 showed pronounced differences for PMNs that barely missed statistical significance (Fig. 3A). As aforementioned, platelet association is not uncommon for leukocytes, especially in the context of inflammatory disorders. Whereas PMN-platelet aggregates would persist in unmanipulated whole blood, Ficoll density gradient purification was shown to separate PMNs and platelets due to their different densities (Chanarat and Chiewsilp, 1975). In order to check whether increased platelet marker expression on PMNs was due to *de-novo* surface expression on PMNs or due to platelet association to PMNs, we analyzed whole blood and Ficoll-purified PMNs together with PBMCs from the same patients and healthy controls that were processed identically. Indeed, compared to whole blood staining, where psoriasis patient samples showed higher CD41 and CD61 expression than healthy donors (Fig. 3B), Ficoll purification reduced these differences in CD41 and CD61 staining between psoriasis patients and healthy donors: Ficoll-purified PMNs showed more similar levels of CD41 and CD61 positivity between the two groups (Fig. 3C). As Ficoll is unlikely to lead to a shedding of CD41 or CD61 (for example, CD62L was unaffected), we concluded that the platelet surface markers seen in psoriasis PMNs in whole blood analysis were due to platelet-PMN conjugate formation. Similar results were obtained for monocytes (Fig. S3A-C). This was confirmed by bright-field fluorescence microscopy which showed small CD41^+^ CD62P^+^ entities, clearly interpretable as platelets, attached to the outer surface of the CD66b^+^ PMNs (Fig. 3D). Including anti-CD62P (P-Selectin) Abs in the analysis, we noted that the platelets were CD62P ^+^ (Fig. 3E) i.e. activated, in line with recent analyses (Sreeramkumar et al., 2014). Thus the expression of platelet markers on psoriasis PMNs is attributable to an increased formation of platelet-PMN aggregates.

To gain deeper insight into whether the formation of PMN- and monocyte-aggregates may be a result of skin inflammation, we analyzed aggregate formation in the imiquimod (IMQ) *in vivo* mouse model of experimental psoriasis. In this model the application of the TLR7 agonists imiquimod leads to a fulminant local and psoriasiform inflammation accompanied by PMN influx (Gilliet et al., 2004). Fig. 3F and G show that IMQ-treated mice exhibited significantly higher PMN-platelet and monocyte-platelet aggregates compared to control mice, in agreement with the data obtained from human psoriasis patients. Collectively, these results indicate that PMN-platelet and monocyte-platelet aggregates are increased in psoriasis and co-incide with skin inflammation.

### Platelets are detectable in psoriatic skin and show distinct surface antigens in the circulation of psoriasis patients

Although platelet activation is known to promote leukocyte association, we wondered whether a mere increase in total platelet counts would correlate with psoriasis as an increased platelet count might favor platelet-PMN association. As expected, platelet counts in the circulation of human psoriasis patients were 62% higher than in healthy donors (p<0.0001, Fig. 4A). To investigate if this higher abundance might also pertain to the skin of patients, where PMNs are frequently infiltrating, we stained for platelet markers in psoriasis skin, which according to our knowledge has not been done before. Whereas neither PMNs (identified by anti-neutrophil elastase, NE, Abs) nor platelets (stained using anti-CD41 and -CD42b) were rarely detectable in healthy skin or isotype-stained samples, in psoriasis skin samples both cell types were present (Fig. 4B, quantified in Fig. 4C). Certain samples even showed co-localization of PMNs and platelets within infiltrated areas (Fig. 4D, quantified in E). This indicates that platelets are only infiltrating lesional skin, and may preferentially do so in tandem with PMNs.

**Figure 4:**
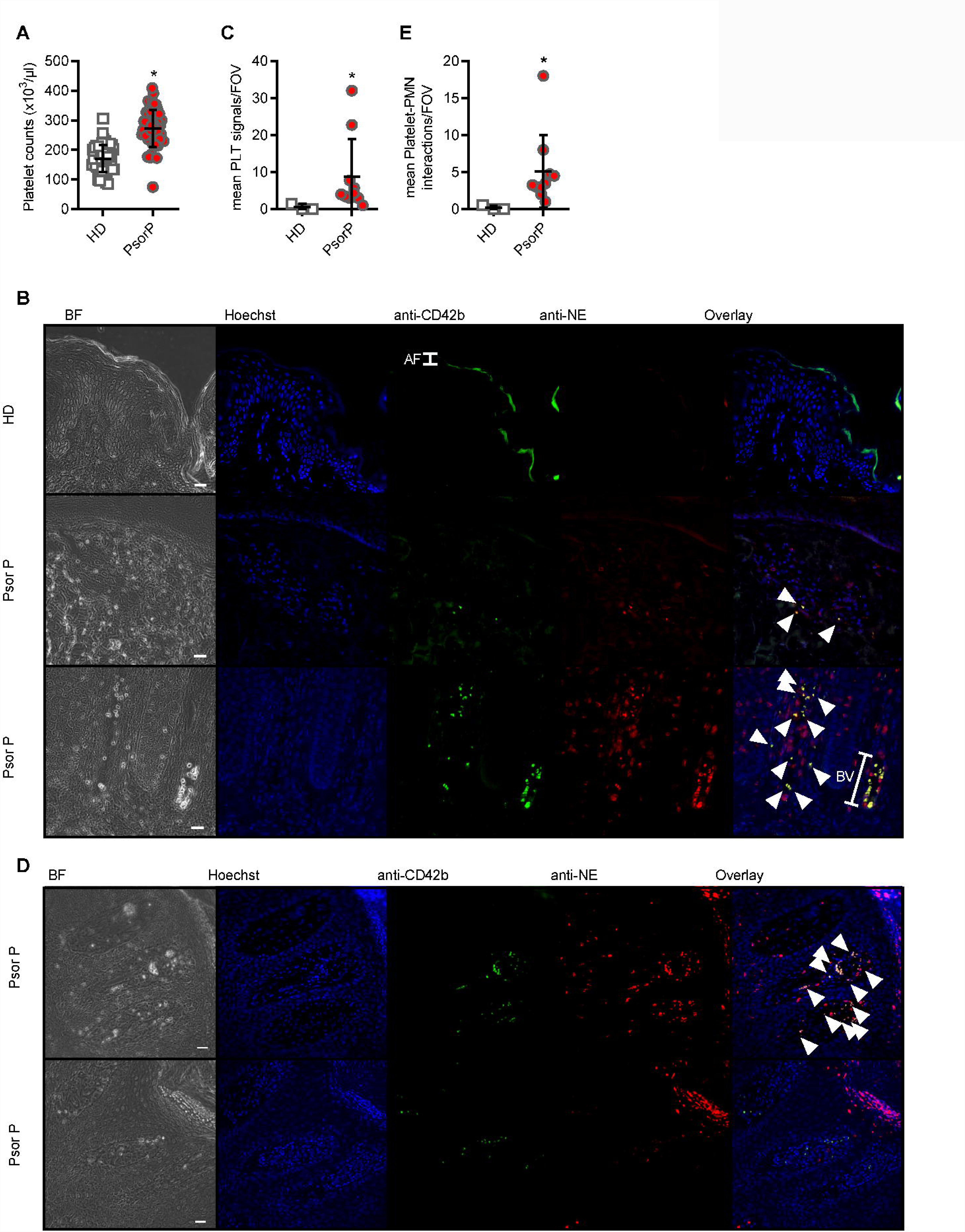
Platelets are more abundant in the blood and skin of psoriasis patients. Total platelet counts in HDs and PsorP (A, n=52 vs 53). (B) Fluorescence microscopy analysis of skin sections stained as indicated from healthy donor or psoriasis-affected skin (n=12 patients and 3 healthy controls, scale bar = 20 μm) and quantified in (C) (D) as in B but showing platelet-PMN aggregates in certain skin samples, quantified in (E). AF = autofluorescence, BV = blood vessel. A, C and E represent combined data (mean+SD) from ‘n’ biological replicates (each dot represents one donor or mouse). In B and D representatives of ‘n’ biological replicates (donors) are shown (mean+SD). * p<0.05 according to an unpaired Student’s t-test (A) or Mann-Whitney test (B, C).

### **Interference with platelets *in vivo* ameliorates skin pathology**

To gain a first impression whether the occurrence of platelet-leukocyte aggregates and the presence of platelets in the psoriatic skin may be causally involved in skin pathology in the context of psoriasis, we investigated the effects of platelet depletion in the IMQ model. Consistently with previous results (Sumida et al., 2014), PMNs proved relevant for disease severity as shown using a PMN-depleting Ab (anti-Ly6G, Fig. 5A). Importantly, in vivo depletion of circulating platelets on d1 and d3 of IMQ-treatment using anti-CD42b antibodies (Elzey et al., 2003) but not control IgG showed a significant decrease in ear skin thickness (Fig. 5B) and epidermal dysplasia (Fig. 5C, quantified in D), concomitant with the verified depletion of total platelets in circulation (Fig. 5E). Consistent with the decrease in PMN-platelet aggregates (Fig. 5F), the number of free PMN was significantly increased in the blood (Fig. 5G). Taken together, these data indicate that circulating platelets significantly contribute to IMQ-induced skin inflammation and pathology.

**Figure 5:**
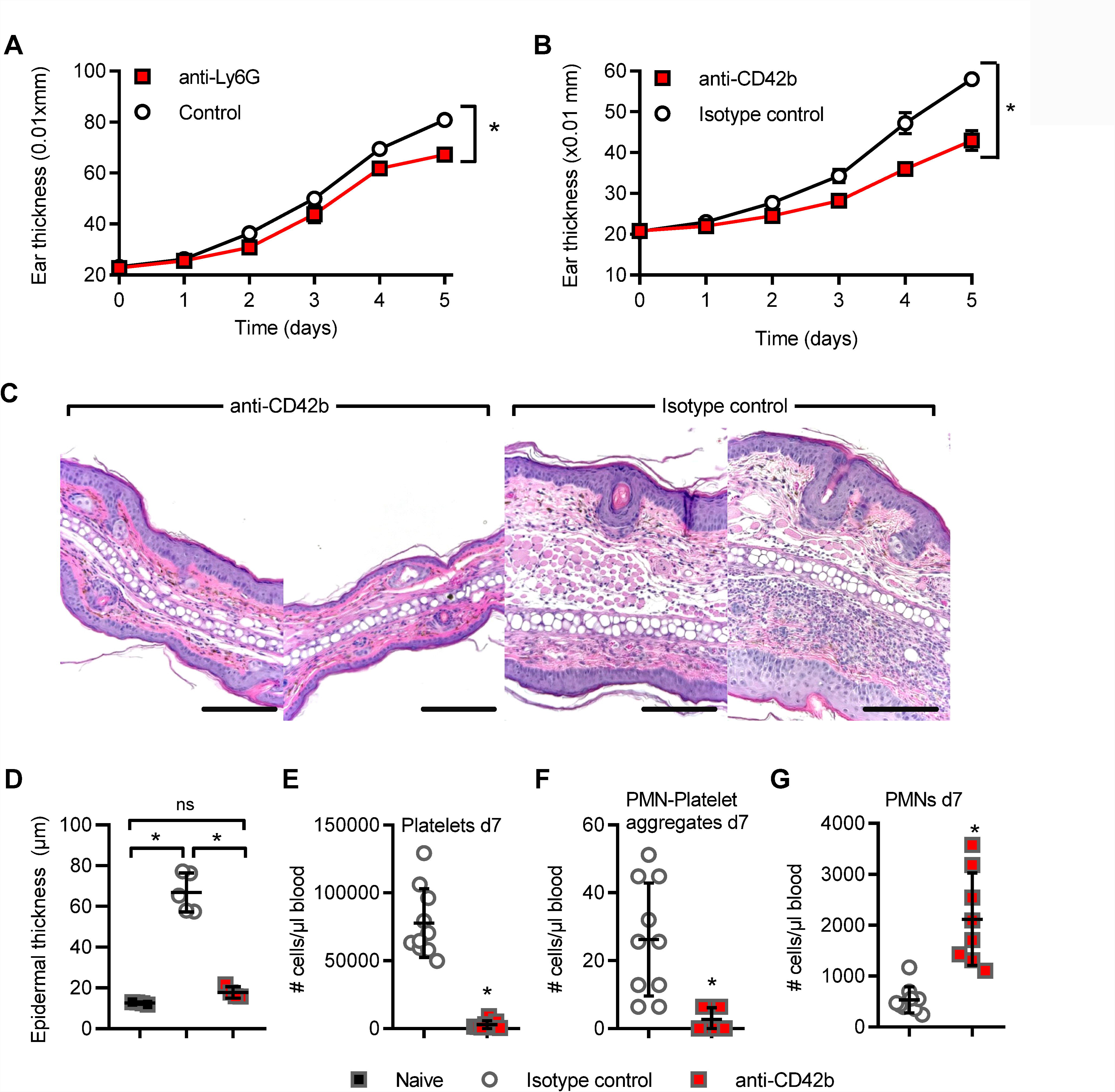
In vivo interference with platelets in experimental psoriasis model. (A-B) Imiquimod-induced experimental psoriasis model in BL/6 mice with ear thickness (mm x 0.01) measured upon PMN depletion by anti-Ly6G (A) or anti-CD42b (B) or respective isotype controls. (C) representative H&E staining from anti-CD41 and isotype-treated skin (n=4-5, scale bar = 180 μm), quantified in D. (E-G) Flow cytometry analysis of total platelet counts on d5 (E), PMN-platelet (F) and free PMNs (G) on d7 upon infusion of IMQ-treated animals with anti-CD42b Abs or isotype control (n=10 each). A, B, D-G represent combined data (mean+SD) from ‘n’ biological replicates (each dot represents one mouse). In C one representative of ‘n’ biological replicates (mouse biopsies) is shown * p<0.05 according to two-way ANOVA (A, B), unpaired Student’s t-test (D), Mann-Whitney test (E-G).

## Discussion

In this present study we screened the surface antigens in different immune cells in the blood of psoriasis patients and identified respective alterations compared to healthy controls, most notably an association of PMNs with platelets. Although the power of our analysis is limited in terms of sample size and only nominal p-values calculated for individual surface antigens, PCA analysis showed different combinations of surface receptors to distinguish patients form healthy donors. It will be informative to analyze the suggested surface antigen combinations in larger cohorts and in the course of different topical or systemic psoriasis treatments. The whole blood-based protocol developed here may be useful for such a later/future application.

Psoriasis has long been viewed as an immunologically-driven disease, in which different leukocytes and T cells play important roles. Although platelet interactions with leukocytes have been reported for psoriasis mouse models and are typically found in many other inflammatory diseases (Ludwig et al., 2004;Tamagawa-Mineoka, 2015;Finsterbusch et al., 2018), a specific role for platelets in the skin lesions of psoriasis patients has not been noted. In agreement with the systemic inflammatory nature of psoriasis, we found platelet-PMN and -monocyte aggregates in the circulation to correlate with the disease in the skin. Of note, the depletion of circulating platelets - and consequently aggregates thereof - unexpectedly and drastically ameliorated skin disease. Our data are in line with platelet depletion experiments conducted in an experimental model of atopic dermatitis, another inflammatory skin disease characterized by severe itching of the skin (Tamagawa-Mineoka et al., 2007). For both atopic dermatitis and psoriasis it could be envisaged that extravasation of leukocytes from skin-proximal blood vessels is facilitated by bound platelets and these platelets then are transported along into the skin. Whether such a scenario of ‘piggyback homing’ applies only to the skin or whether other organs apart from the skin are also affected remains to be investigated but may unearth a possible link to the cardiovascular comorbidities observed in psoriasis patients. It is also conceivable that platelets that thus end up in the skin could directly contribute to disease severity - either in concert with leukocytes (Li et al., 2015; lba and Levy, 2018) or acting separately. The responsiveness of platelets to microbe- and danger-associated molecular patterns, ability for active locomotion and for secretion of inflammatory mediators described for platelets may be properties that could be involved (Tamagawa-Mineoka, 2015);(Semple et al., 2011). Although a feedback to myelopoiesis can also not be excluded, the increase of free PMNs in the blood of platelet-depleted mice would on the other hand suggest that the absence of circulating platelets traps PMNs in circulation and prevents skin pathology by preventing PMN skin infiltration. If so, this would add psoriasis to the list of pathologies in which therapeutics preventing platelet-PMN interactions may prove beneficial but possibly at the expense of reducing peripheral innate defenses which rely on PMNs being able to enter the periphery (Sreeramkumar et al., 2014).

Platelets have been credited with ability to promote peripheral homing of innate immune cells. However, the observation that platelets themselves enter psoriatic skin was unexpected. Further exploration of the role of skin-infiltrating platelets and their mechanisms of skin-homing or even active migration (Gaertner et al., 2017) in psoriasis and other inflammatory conditions may thus prove informative. On the other hand, the role of platelets in the cardiovascular comorbidities often associated with psoriasis is intriguing. Given the sensitivity of platelets to IL-17 (Maione et al., 2011), a key psoriasis-associated cytokine, investigation of the IL-17-axis and effects of anti-IL-17 biologicals on isolated platelets and platelet-leukocyte aggregates may prove insightful in this context.

## Acknowledgements

We thank S. Kohler and B. Franklin for technical support and helpful discussions and all healthy donors and patients for participation in our study. This study was supported by the Deutsche Forschungsgemeinschaft (German Research Foundation, DFG) CRC TR156 “The skin as an immune sensor and effector organ - Orchestrating local and systemic immunity”, the University of Tübingen and the University Hospital Tübingen.

## Authorship contributions

FH, ZB, NA performed experiments; ANRW, FH, MCC, NA, SH analyzed data; MH, MWL, JW, RB, MS, DS, KS, SG, KG, LM were involved in sample and reagent acquisition; FH and ANWR were involved in the conceptual development of the study, wrote the manuscript and all authors commented on or revised the manuscript; ANRW coordinated and supervised the entire study.

## Abbreviations

MFI: Mean fluorescence intensity
MAMPs: Microbe-associated molecular pattern
IMQ: Imiquimod
IFN: Interferon
IL: Interleukin
KC: Keratinocytes
PLT: Platelet
PMNs: Polymorphonuclear neutrophils
PCA: Principal component analysis

## References

Bates, D., Mächler, M., Bolker, B., and Walker, S. (2015). Fitting Linear Mixed-Effects Models Using Ime4. 2015 67, 48.

Bates, J.C.P.D.M., Pinheiro, J., Pinheiro, J.C., and Bates, D. (2000). Mixed-Effects Models in S and S-PLUS. Springer New York.

Boehncke, W.H. (2018). Systemic Inflammation and Cardiovascular Comorbidity in Psoriasis Patients: Causes and Consequences. Front Immunol 9, 579.

Bretz, F., Hothorn, T., and Westfall, P.H. Multiple comparisons using R.

Caiazzo, G., Fabbrocini, G., Di Caprio, R., Raimondo, A., Scala, E., Balato, N., and Balato, A. (2018). Psoriasis, Cardiovascular Events, and Biologies: Lights and Shadows. Front Immunol 9,1668.

Chanarat, P., and Chiewsilp, P. (1975). A simple method for the elimination of platelets from the lymphocyte-platelet mixture by sucrose. Am J Clin Pathol 63, 237–239.

De Bruijne-Admiraal, L.G., Modderman, P.W., Von Dem Borne, A.E., and Sonnenberg, A. (1992). P-selectin mediates Ca(2+)-dependent adhesion of activated platelets to many different types of leukocytes: detection by flow cytometry. Blood 80,134–142.

Eberle, F.C., Bruck, J., Holstein, J., Hirahara, K., and Ghoreschi, K. (2016). Recent advances in understanding psoriasis. FlOOORes 5.

Elzey, B.D., Tian, J., Jensen, R.J., Swanson, A.K., Lees, J.R., Lentz, S.R., Stein, C.S., Nieswandt, B., Wang, Y., Davidson, B.L., and Ratliff, T.L. (2003). Platelet-mediated modulation of adaptive immunity. A communication link between innate and adaptive immune compartments. Immunity 19, 9–19.

Finsterbusch, M., Schrottmaier, W.C., Kral-Pointner, J.B., Salzmann, M., and Assinger, A. (2018). Measuring and interpreting platelet-leukocyte aggregates. Platelets 29, 677–685.

Gaertner, F., Ahmad, Z., Rosenberger, G., Fan, S., Nicolai, L., Busch, B., Yavuz, G., Luckner, M., Ishikawa-Ankerhold, H., Hennel, R., Benechet, A., Lorenz, M., Chandraratne, S., Schubert, I., Helmer, S., Striednig, B., Stark, K., Janko, M., Bottcher, R.T., Verschoor, A., Leon, C., Gachet, C., Gudermann, T., Mederos, Y.S.M., Pincus, Z., Iannacone, M., Haas, R., Wanner, G., Lauber, K., Sixt, M., and Massberg, S. (2017). Migrating Platelets Are Mechano-scavengers that Collect and Bundle Bacteria. Cell 171,1368–1382 el323.

Gilliet, M., Conrad, C., Geiges, M., Cozzio, A., Thurlimann, W., Burg, G., Nestle, F.O., and Dummer, R. (2004). Psoriasis triggered by toll-like receptor 7 agonist imiquimod in the presence of dermal plasmacytoid dendritic cell precursors. Arch Dermatol 140,1490–1495.

Golden, J.B., Groft, S.G., Squeri, M.V., Debanne, S.M., Ward, N.L., Mccormick, T.S., and Cooper, K.D. (2015). Chronic Psoriatic Skin Inflammation Leads to Increased Monocyte Adhesion and Aggregation. J Immunol 195, 2006–2018.

Graessel, A., Hauck, S.M., Von Toerne, C., Kloppmann, E., Goldberg, T., Koppensteiner, H., Schindler, M., Knapp, B., Krause, L., Dietz, K., Schmidt-Weber, C.B., and Suttner, K. (2015). A Combined Omics Approach to Generate the Surface Atlas of Human Naive CD4+ T Cells during Early T-Cell Receptor Activation. Mol Cell Proteomics 14, 2085–2102.

Habets, K.L., Huizinga, T.W., and Toes, R.E. (2013). Platelets and autoimmunity. Eur J Clin Invest 43, 746–757.

Iba, T., and Levy, J.H. (2018). Inflammation and thrombosis: roles of neutrophils, platelets and endothelial cells and their interactions in thrombus formation during sepsis. J Thromb Haemost 16, 231–241.

Jungi, T.W., Spycher, M.O., Nydegger, U.E., and Barandun, S. (1986). Platelet-leukocyte interaction: selective binding of thrombin-stimulated platelets to human monocytes, polymorphonuclear leukocytes, and related cell lines. Blood 67, 629–636.

Langewouters, A.M.G., Van Erp, P.E.J., De Jong, E.M.G.J., and De Kerkhof, P.C.M.V. (2008). Lymphocyte subsets in peripheral blood of patients with moderate-to-severe versus mild plaque psoriasis. Archives of Dermatological Research 300,107–113.

Lenth, R.V. (2016). Least-Squares Means: The R Package lsmeans. Journal of Statistical Software 69, 1–33.

Li, J., Kim, K., Barazia, A., Tseng, A., and Cho, J. (2015). Platelet-neutrophil interactions under thromboinflammatory conditions. Cell Mol Life Sci 72, 2627–2643.

Ludwig, R.J., Schultz, J.E., Boehncke, W.H., Podda, M., Tandi, C., Krombach, F., Baatz, H., Kaufmann, R., Von Andrian, U.H., and Zollner, T.M. (2004). Activated, not resting, platelets increase leukocyte rolling in murine skin utilizing a distinct set of adhesion molecules. J Invest Dermatol 122, 830–836.

Maione, F., Cicala, C., Liverani, E., Mascolo, N., Perretti, M., and D’acquisto, F. (2011). IL-17A increases ADP-induced platelet aggregation. Biochem Biophys Res Commun 408, 658–662.

Schindelin, J., Arganda-Carreras, I., Frise, E., Kaynig, V., Longair, M., Pietzsch, T., Preibisch, S., Rueden, C., Saalfeld, S., Schmid, B., Tinevez, J.Y., White, D.J., Hartenstein, V., Eliceiri, K., Tomancak, P., and Cardona, A. (2012). Fiji: an open-source platform for biological-image analysis. Nat Methods 9, 676–682.

Schon, M.P., Broekaert, S.M., and Erpenbeck, L. (2017). Sexy again: the renaissance of neutrophils in psoriasis. Exp Dermatol 26, 305–311.

Schon, M.P., and Erpenbeck, L. (2018). The Interleukin-23/lnterleukin-17 Axis Links Adaptive and Innate Immunity in Psoriasis. Front Immunol 9,1323.

Schuler, R., Brand, A., Klebow, S., Wild, J., Protasio Veras, F., Ullmann, E., Roohani, S., Kolbinger, F., Kossmann, S., Wohn, C., Daiber, A., Munzel, T., Wenzel, P., Waisman, A., Clausen, B.E., and Karbach, S. (2018). Antagonization of IL-17A attenuates skin inflammation and vascular dysfunction in mouse models of psoriasis. J Invest Dermatol.

Semple, J.W., Italiano, J.E., Jr., and Freedman, J. (2011). Platelets and the immune continuum. Nat Rev Immunol 11, 264–274.

Sen, B.B., Rifaioglu, E.N., Ekiz, O., Inan, M.U., Sen, T., and Sen, N. (2013). Neutrophil to lymphocyte ratio as a measure of systemic inflammation in psoriasis. Cutaneous and ocular toxicology.

Sreeramkumar, V., Adrover, J.M., Ballesteros, I., Cuartero, M.I., Rossaint, J., Bilbao, I., Nacher, M., Pitaval, C., Radovanovic, I., Fukui, Y., Mcever, R.P., Filippi, M.D., Lizasoain, I., Ruiz-Cabello, J., Zarbock, A., Moro, M.A., and Hidalgo, A. (2014). Neutrophils scan for activated platelets to initiate inflammation. Science 346,1234–1238.

Sumida, H., Yanagida, K., Kita, Y., Abe, J., Matsushima, K., Nakamura, M., Ishii, S., Sato, S., and Shimizu, T. (2014). Interplay between CXCR2 and BLT1 facilitates neutrophil infiltration and resultant keratinocyte activation in a murine model of imiquimod-induced psoriasis. J Immunol 192, 4361–4369.

Tamagawa-Mineoka, R. (2015). Important roles of platelets as immune cells in the skin. J Dermatol Sci 77,93–101.

Tamagawa-Mineoka, R., Katoh, N., Ueda, E., Takenaka, H., Kita, M., and Kishimoto, S. (2007). The role of platelets in leukocyte recruitment in chronic contact hypersensitivity induced by repeated elicitation. Am J Pathol 170, 2019–2029.

Vena, G.A., Vestita, M., and Cassano, N. (2010). Psoriasis and cardiovascular disease. Dermatol Ther 23,144–151.

Zuchtriegel, G., Uhl, B., Puhr-Westerheide, D., Pornbacher, M., Lauber, K., Krombach, F., and Reichel, C.A. (2016). Platelets Guide Leukocytes to Their Sites of Extravasation. PLoS Biol 14, el002459.

