## Supplementary Information text for "Platelet-neutrophil aggregates promote skin pathology in psoriasis"

**Title**

**Running title**

Platelet-neutrophil aggregates in psoriasis

**Authors**

Franziska Herster^1^, Zsofia Bittner^1^, Marius Cosmin Cordrea^2^, Nate Archer^3^, Martin Heister^4^, Markus W. Löffler^1,5^, Simon Heumos^2^, Joanna Wegner^6^, Ramona Businger^7^, Michael Schindler^7^, David Stegner^8^, Knut Schäkel^9^, Stephan Grabbe^6^, Kamran Ghoreschi^4,10^, Lloyd Miller^4^, Alexander N.R. Weber^1,^*

### Supplementary Materials and Methods

##### Reagents

All chemicals were from Sigma unless otherwise stated. Antibodies used for flow cytometry and microscopy are listed in Table S4.

##### Mice

C57Bl/6J mice were purchased from Jackson Laboratories (Bar Harbor, ME). Mice were bred and maintained under the same specific pathogen-free conditions at an American Association for the Accreditation of Laboratory Animal Care (AAALAC)-accredited animal facility at Johns Hopkins University and handled according to procedures described in the Guide for the Care and Use of Laboratory Animals as well as Johns Hopkins University’s policies and procedures as set forth in the Johns Hopkins University Animal Care and Use Training Manual, and all animal experiments were approved by the Johns Hopkins University Animal Care and Use Committee. Gender-and age-matched 6-8 week old mice were used for each experiment.

##### Study participants and sample acquisition

All patients and healthy blood donors included in this study provided their written informed consent before study participation. Approval for use of their biomaterials was obtained by the local ethics committee at the University of Tübingen, in accordance with the principles laid down in the Declaration of Helsinki as well as applicable laws and regulations. Whole blood from voluntary healthy donors was obtained at the University of Tübingen, Department of Immunology. Blood samples from psoriasis patients were obtained at the University Hospital Tübingen, Department of Dermatology. Psoriasis patients had a median age of median age 41.8 years with PASI scores ≥10 (except for 2 donors in Figs. 3B,C and S3B,C where PASI was >4.5) and did not receive any systemic treatments at the time of blood sampling. All samples obtained from psoriasis patients were processed simultaneously with samples from at least one healthy donor matched for age and sex.

##### Fluorescence Microscopy

200 µl of cell suspension after erythrocyte lysis was used per well (96 well plate). FcR block, staining, fixation and permeabilization was performed as for Flow cytometry. The cell pellets were resuspended in 50-100 µl FACS buffer. 40 µl of the cell suspension was pipetted on a Poly-L-Lysine coated coverslip (Corning, 734-1005) and the cells were left to attach for one hour in the dark. ProLong Diamond Antifade (Life technologies, P36965) was used to mount the coverslips on uncoated microscopy slides. The slides were left to dry overnight at RT in the dark and were then stored at 4 °C before microscopy. The measurements were conducted with a Nikon Ti2 eclipse (100 x magnifications) and the analysis was performed using ImageJ/Fiji analysis software.

##### Imiquimod model of psoriatic skin inflammation

Mice were anesthetized (2% isoflurane) and 62.5mg of 5% imiquimod (Tora Pharmaceuticals) was applied topically with a sterile cotton swab to the ventral and dorsal side of the mouse ear daily for a total of 5 treatments. Prior to imiquimod application, ear thickness was measured with a manual caliper (Peacock). A day before and at the end of imiquimod treatment, retro-orbital blood samples were collected with heparinized capillary tubes (Fisher) for FACS analysis. In addition, full thickness ear skin was excised with a 6mm punch (Acuderm) for histological analysis.

##### Platelet depletion protocol

For platelet depletion, 4 µg/g of anti-CD42b (Emfret Analytics, R300) or rat IgG isotype control (Emfret Analytics, R301) in sterile PBS was administered i.v. one day before, and 2 µg/g administered i.p. 3 days after, the first imiquimod treatment.

##### Flow cytometry

A 50 µl sample of blood was first diluted in 300 µl TBS containing 5 U/ml Heparin followed by a dilution with 500 µl PBS. A volume of 5 µl of diluted sample was stained with the following mAbs for FACS analysis: anti-Ly6C, anti-CD45, anti-CD41, anti-Ly6G and anti-CD11b. Propidium iodide was used to measure cell viability and TruStain fcX was used to block Fc receptor binding.

##### Histology and epidermal thickness measurements

6-mm punch biopsy specimens were placed in 10% formalin and paraffin-embedded. Skin cross-sections (4 μm) were prepared and stained with hematoxylin-eosin (H&E) by the Johns Hopkins Reference Histology Laboratory according to clinical specimen guidelines, or utilized for immunofluorescent staining. To measure epidermal thickness, at least 10 epidermal thickness measurements per mouse were averaged from images taken at 20x magnification (Leica, DFC495 or ECHO Revolve) using ImageJ/Fiji software.

##### Statistics

Experimental data was analyzed using Excel 2010 (Microsoft) and/or GraphPad Prism 6, 7 or 8 (GraphPad Software, Inc.), microscopy data with ImageJ/Fiji, flow cytometry data using FlowJo software version 10. When extreme values occurred, outliers were statistically identified using the ROUT test at high (0.5%) stringency and normality tested using the Shapiro-Wilk test for the subsequent choice of a parametric or non-parametric tests. p-values (α=0.05, β=0.8) were then calculated and multiple testing was corrected for in Prism as indicated in figure legends. Values < 0.05 generally considered statistically significant. These were denoted by * throughout, even if the calculated p-values were considerably lower than 0.05. Comparisons made to unstimulated control unless indicated otherwise by brackets.

### Supplementary Figure Legends

#### Supplementary figure S1

Gating strategy used in the surface antigen screen in human whole blood. (B-E) Selected surface antigens with significant differences in MFI between healthy donors (HDs) and psoriasis patients (PsorP), n=5 each based on a representative dataset.

#### Supplementary figure S2

(A-C) Selected additional surface antigens with significant differences in MFI between the PMNs (A), monocytes (B) and B cells (C) from healthy donors (HDs) and psoriasis patients (PsorP), n=5 each. A-C represent combined data (mean+SD) from ‘n’ biological replicates. * p<0.1 nominal by two-way ANOVA followed by Tukey’s multiple comparisons correction.

#### Supplementary figure S3

(A) Expression analysis screening results for CD41 (upper) and CD61 (lower panel) MFIs on gated monocytes between healthy donors (HDs) and psoriasis patients (PsorP), n=5 each. (B+C) Flow cytometric analysis of CD41- or CD61-positive gated monocytes in HD or PsorP (HD n=7, PsorP n=5) in whole blood samples (B) or Ficoll density gradient centrifugation. A-C represent combined data (mean+SD) from ‘n’ biological replicates (each dot represents one donor). * p<0.1 nominal by two-way ANOVA followed by Tukey’s multiple comparisons correction, in B and C * p<0.05 by unpaired Student’s t-Tests.

#### Supplementary figure S4

Selected additional surface antigens with significant differences in MFI between platelets from healthy donors (HDs) and psoriasis patients (PsorP), n=5 each. A represents combined data (mean+SD) from ‘n’ biological replicates. * p<0.1 nominal by two-way ANOVA followed by Tukey’s multiple comparisons correction.

### Supplementary Tables

**Table S1: Antibodies**

| Item | Fluorophore | Species | Isotype | Company | Product no. |
| --- | --- | --- | --- | --- | --- |
| Isotype control | PE | mouse | IgG1 kappa | eBioscience | 12471442 |
| Isotype control | FITC | mouse | IgM | BioLegend | 401605 |
| Isotype control | BV421 | mouse | IgG1 kappa | BioLegend | 400157 |
| Isotype control | AF488 | mouse | IgG2b kappa | BioLegend | 402207 |
| Isotype control | AF488 | mouse | IgG1 | BioLegend | 400134 |
| Isotype control | AF647 | mouse | IgM | BioLegend | 401618 |
| Isotype control | PE-Cy7 | mouse | IgG1 | BioLegend | 400126 |
| Anti-hCD15 | PE | mouse | IgG1 kappa | BioLegend | 323006 |
| Anti-hCD66b | FITC | mouse | IgG1 kappa | BioLegend | 305103 |
| Anti-hCD62L | BV421 | mouse | IgG1 kappa | BioLegend | 30482 |
| Anti-hCD3 | AF488 | mouse | IgG1 kappa | BioLegend | 317310 |
| Anti-hCD19 | BV421 | mouse | IgG1 | BioLegend | 302234 |
| Anti-hCD15 | PE-Cy7 | mouse | IgG1 kappa | BioLegend | 323030 |
| Zombie yellow fixable dye | - | - | - | BioLegend | 423103 |
| Anti-hCD41 | PE | mouse | IgG1 | BioLegend | 303706 |
| Anti-hCD61 | PE | mouse | IgG1 | BioLegend | 336406 |
| Anti-hCD66b | AF647 | mouse | IgM | BioLegend | 305109 |
| Anti-hCD62P | AF488 | mouse | IgG1 | BioLegend | 304916 |
| Anti-hCD41 | unconjugated | rabbit | IgG1 | Abcam | ab63983 |
| Anti-hCD42b | unconjugated | goat | IgG | Santa Cruz | sc-7070 |
| Anti-hNeutrophil Elastase | unconjugated | mouse | IgG1 | Novus Biologicals | MAB91671-100 |
| Anti-rabbit IgG | AF647 | chicken | IgY | ThermoFisher | A-21443 |
| Anti-mouse IgG1 | AF594 | chicken | IgY | ThermoFisher | A-21201 |
| Anti-goat IgG | AF488 | chicken | IgY | ThermoFisher | A-21467 |
| anti-mouse Ly6C | VioBlue | rat | IgG2a | Miltenyi | 130-102-929 |
| Anti-mouse CD45 | APC-Vio770 | rat | IgG2bκ | Miltenyi | 130-118-687 |
| Anti-mouse CD41 | FITC | rat | IgG1κ | Miltenyi | 130-105-929 |
| Anti-mouse Ly6G | PE | rat | IgG1κ | Miltenyi | 130-102-895 |
| Anti-mouse CD11b | APC | rat | IgG2bκ | Miltenyi | 130-113-793 |
| Propidium iodide | - | - | - | Miltenyi | 130-093-233 |
| TruStain fcX | - | - | - | BioLegend | 422302 |

#### Table S2: 10x Ammonium chloride erythrocyte lysis buffer

| Chemical | Company | Product no. |
| --- | --- | --- |
| 1.54 M NH_4_Cl | Roth | 5470.1 |
| 100 mM KHCO_3_ | Fluka | 60220 |
| 1 mM EDTA; pH 8 | ThermoFisher | 15575020 |
| dissolved in Ampuwa water | Fresenius Kabi | 1833 |
| pH adjusted to 7.3, sterile filtered (0.22 µm) |  |  |

Bates, D., Mächler, M., Bolker, B., and Walker, S. (2015). Fitting Linear Mixed-Effects Models Using lme4. *2015* 67**,** 48.

Bates, J.C.P.D.M., Pinheiro, J., Pinheiro, J.C., and Bates, D. (2000). *Mixed-Effects Models in S and S-PLUS.* Springer New York.

Boehncke, W.H. (2018). Systemic Inflammation and Cardiovascular Comorbidity in Psoriasis Patients: Causes and Consequences. *Front Immunol* 9**,** 579.

Bretz, F., Hothorn, T., and Westfall, P.H. *Multiple comparisons using R.*

Caiazzo, G., Fabbrocini, G., Di Caprio, R., Raimondo, A., Scala, E., Balato, N., and Balato, A. (2018). Psoriasis, Cardiovascular Events, and Biologics: Lights and Shadows. *Front Immunol* 9**,** 1668.

Chanarat, P., and Chiewsilp, P. (1975). A simple method for the elimination of platelets from the lymphocyte-platelet mixture by sucrose. *Am J Clin Pathol* 63**,** 237-239.

De Bruijne-Admiraal, L.G., Modderman, P.W., Von Dem Borne, A.E., and Sonnenberg, A. (1992). P-selectin mediates Ca(2+)-dependent adhesion of activated platelets to many different types of leukocytes: detection by flow cytometry. *Blood* 80**,** 134-142.

Eberle, F.C., Bruck, J., Holstein, J., Hirahara, K., and Ghoreschi, K. (2016). Recent advances in understanding psoriasis. *F1000Res* 5.

Elzey, B.D., Tian, J., Jensen, R.J., Swanson, A.K., Lees, J.R., Lentz, S.R., Stein, C.S., Nieswandt, B., Wang, Y., Davidson, B.L., and Ratliff, T.L. (2003). Platelet-mediated modulation of adaptive immunity. A communication link between innate and adaptive immune compartments. *Immunity* 19**,** 9-19.

Finsterbusch, M., Schrottmaier, W.C., Kral-Pointner, J.B., Salzmann, M., and Assinger, A. (2018). Measuring and interpreting platelet-leukocyte aggregates. *Platelets* 29**,** 677-685.

Gaertner, F., Ahmad, Z., Rosenberger, G., Fan, S., Nicolai, L., Busch, B., Yavuz, G., Luckner, M., Ishikawa-Ankerhold, H., Hennel, R., Benechet, A., Lorenz, M., Chandraratne, S., Schubert, I., Helmer, S., Striednig, B., Stark, K., Janko, M., Bottcher, R.T., Verschoor, A., Leon, C., Gachet, C., Gudermann, T., Mederos, Y.S.M., Pincus, Z., Iannacone, M., Haas, R., Wanner, G., Lauber, K., Sixt, M., and Massberg, S. (2017). Migrating Platelets Are Mechano-scavengers that Collect and Bundle Bacteria. *Cell* 171**,** 1368-1382 e1323.
