## Supplementary figures and images for "Platelet-neutrophil aggregates promote skin pathology in psoriasis"

### Figure S1

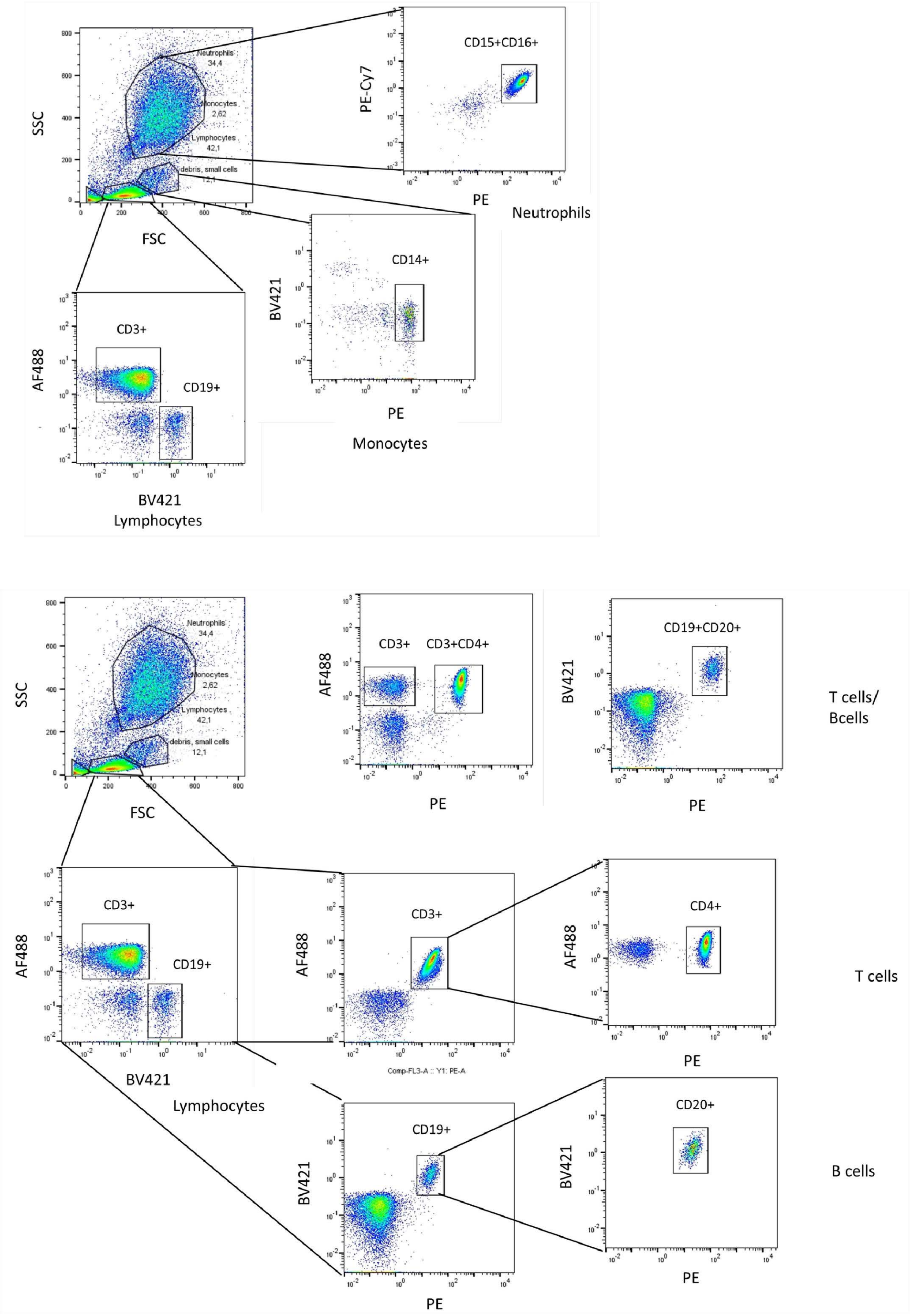

### Figure S2

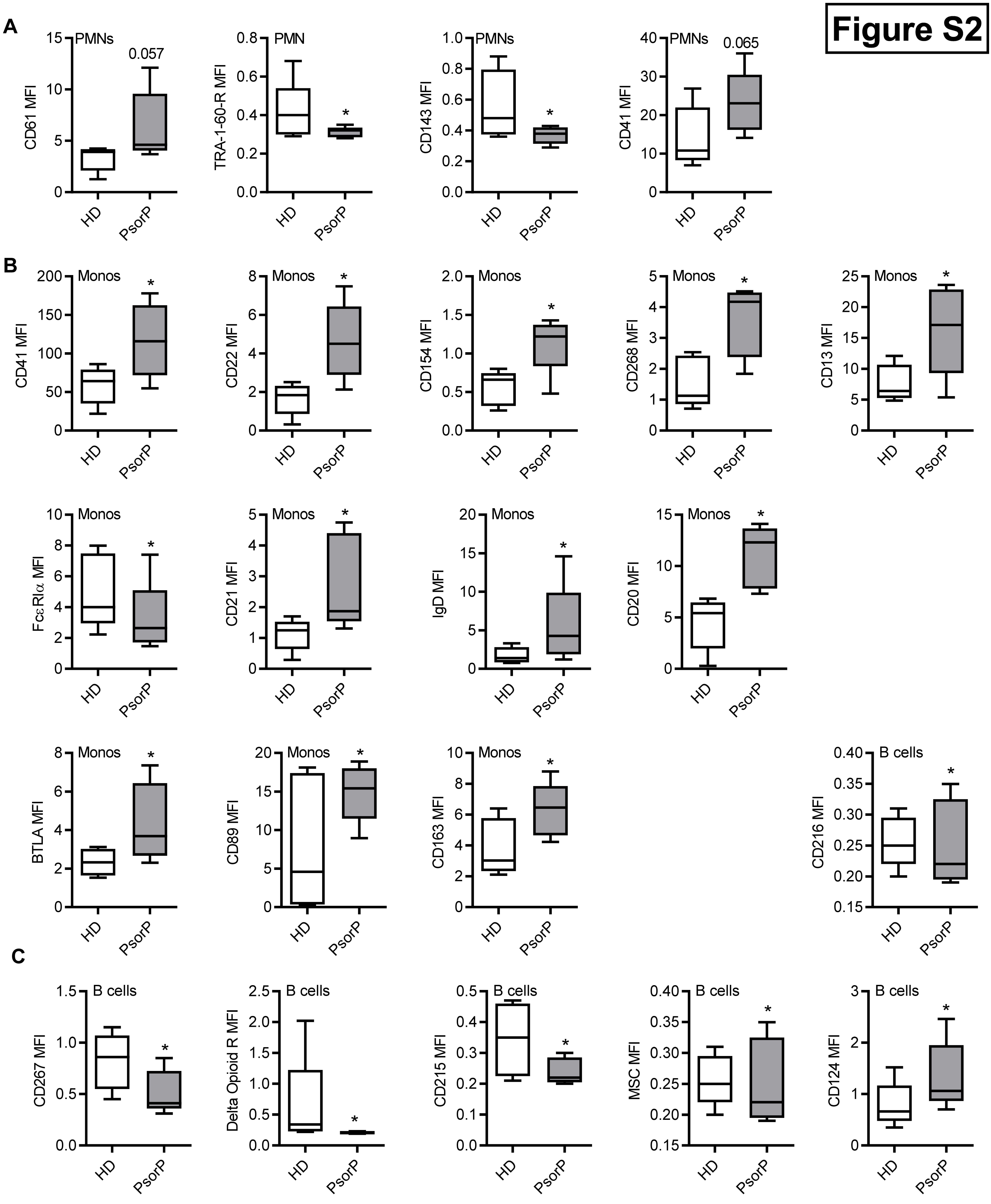

### Figure S3

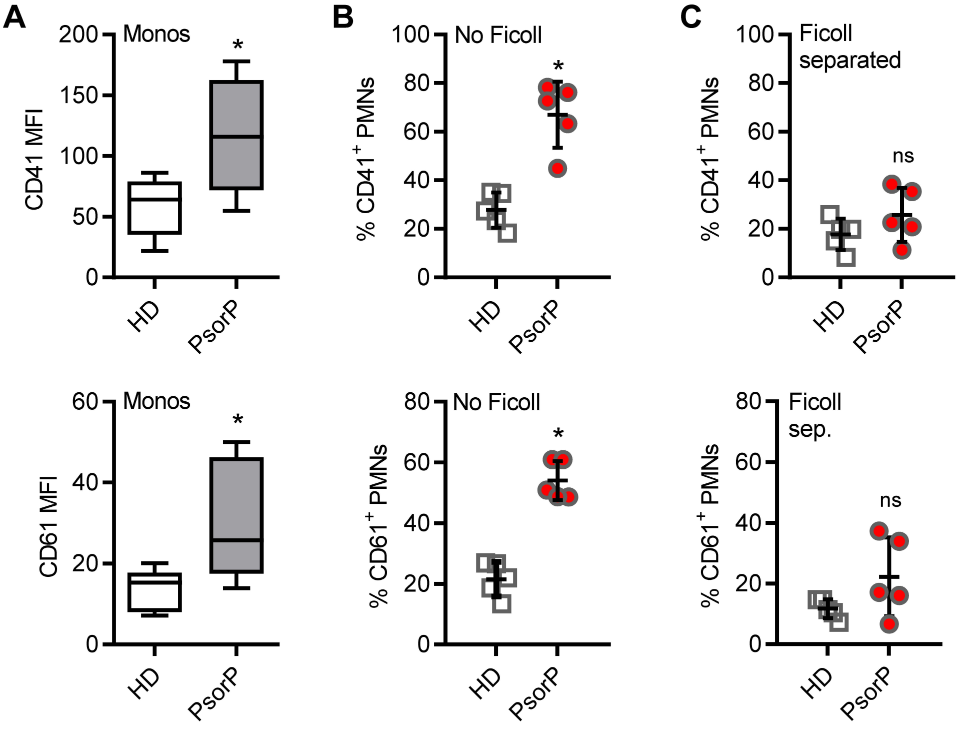
